# Inhibition of NSD1 by 5-O-Sulfamoyl Adenosine improved 5-FU sensitivity by suppressing cancer cell proliferation and xenograft tumor growth

**DOI:** 10.64898/2026.06.10.731397

**Authors:** Zahid Rafiq, Kulbhushan Tikoo

## Abstract

Epigenetics regulate cell-cycle kinetics, differentiation, apoptosis, and migration. Nuclear receptor-binding SET Domain (NSD) histone methyltransferases represent a family of oncoproteins with aberrant expression in cancer. Emerging reports suggest that NSD1 could be an attractive target as its expression is correlated with poor prognosis and tumorigenesis. Previously, we reported the target validation and structure-based virtual screening against NSD1, leading to the selection of several hit molecules with relatively high docking and MMGBSA delta G Bind scores. One of the best-fit molecules identified was 5’-O-sulfamoyl adenosine (5-SA) and was compared with the S-Adenosyl-l-Cysteine (SAC), a structural analog of S-Adenosyl-l-Methionine (SAM) for its inhibitory activity against NSD1. IC_50_ values for 5-SA and SAC against NSD1 were 53.819 µM and 115.003 µM respectively. 5-SA significantly reduced the viability of DU145 and HepG2 cells with IC_50_ values calculated as 198µM and 168.3 µM respectively. It also reduced the RNA and protein expression levels of NSD1 and subsequently prevented dimethylation of lysine 36 on histone H3 (H3K36me2). Furthermore, 5-SA impeded proliferation, and migration, altered the cell cycle phase, and induced cell apoptosis. Interestingly, 5-SA potentiated the anticancer activity of 5-Fluorouracil (5-FU) against cancer cells. The xenograft model of prostate cancer also showed that 5-SA significantly reduced the tumor growth kinetics. However, the combination of 5-SA and 5-FU synergistically reduced tumor growth and improved survival of animals. To the best of our knowledge, we report for the first time that 5-SA mediated inhibition of NSD1 enhanced the tumor sensitivity to 5-FU and thereby, improved the tumor growth and progression.

**Graphical Abstract:** 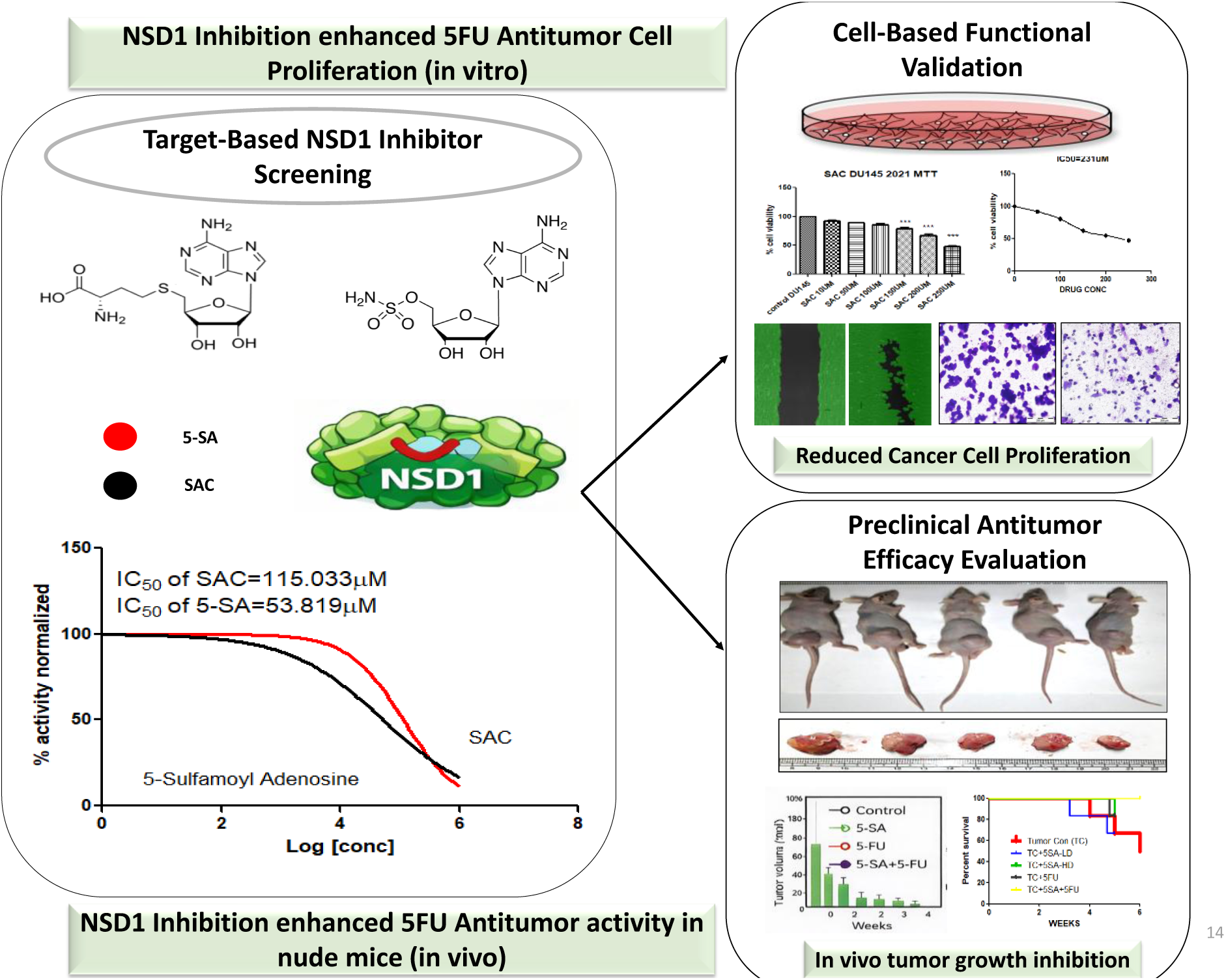

## Introduction

Epigenetic dysregulation of mechanisms including DNA methylation, histone modifications, and microRNA and machinery in the form of readers, writers, and erasers may lead to the process of oncogenesis and other diseases due to aberrant gene expression and function [1,2]. Currently, epigenetic research focuses on the identification of novel targets and the development of potent inhibitors as it represents a very promising, simple, and affordable approach to the otherwise very difficult-to-treat cancers and associated metastasis by gene therapy [3]. NSDs are primarily known to regulate gene expression through methylation of lysine 36 at histone H3 (H3K36) [3–5]. Methylation at histone H3 lysine 36 (H3K36) is a conserved epigenetic mark regulating gene transcription, alternative splicing, and DNA repair [6]. Genes encoding H3K36 methyltransferases (and other KMTases) are commonly over-expressed, mutated, or involved in chromosomal translocations in cancer [6]. Of these KMTases, the NSD proteins are perhaps the best-characterized oncoproteins, as they play important roles in multiple cancer types [7,8]. Accumulating evidence validates the roles of NSD1/NSD2/ NSD3 in tumorigenesis and their special value for epigenetic cancer therapy [9]. Structural studies of the catalytic SET domain of H3KKMTases paved the way and opportunities for the design of small molecule inhibitors [10]. Most importantly, potent inhibitors for most H3K36 KMTases have not yet been developed, justifying the opportunity and challenges associated with this target class.

Recent research highlighted that the NSD1 supports NSD2 in fueling the androgen receptor signaling in prostate cancer [11]. We have recently reported the target validation and selection of top hit molecules against NSD1 through virtual screening and molecular dynamic simulation [12]. NSD1 is associated with acute myeloid leukemia [8,13], multiple myeloma [9], lung cancer [14], metastatic prostate cancer [15], metastatic pancreatic duct adenocarcinoma (PDA) [16] and hepatocellular carcinoma [17]. Selective inhibition of the NSD1 is urgently pursued, as it represents a novel therapeutic hope for cancers with a poor prognosis [18,19]. NSD1 is correlated with high expression in higher stages of cancers and lack of potent inhibitors [20]. This study attempts to provide a novel platform by introducing a virtually and enzymatically validated ligand molecule capable of translating anticancer efficacy *in vitro* and *in vivo* environments and delivering a new approach for enhanced chemosensitivity. In metastatic prostate cancer samples, NSD1 is over-expressed and correlates directly with prostate cancer progression as an epigenetic signature capable of discriminating between malignant and nonmalignant prostate tissue [15]. Moreover, increased NSD1 expression is significantly associated with higher clinical stages, suggesting that NSD1 may be a useful prognostic marker in metastatic pancreatic duct adenocarcinoma [16]. 5-O-sulfamoyladenosine (NSC 133114) has been reported to have anti-leukemia activity [21]. 5-O-sulfamoyl Purines including NSC 133114 (5-O-sulfamoyladenosine) and NSC 750854 (6-desamino derivative of NSC 133114) were reported to be active in both *in vitro* and *in vivo* conditions [22]. However, they have not been properly explored. NSC 750854 was also reported to possess anticancer activity against a range of cell lines *in vitro* and displayed activity against several subcutaneous xenograft studies [22]. The fluoropyrimidine 5-fluorouracil (5-FU) is an antimetabolite drug that is widely used for the treatment of several cancers including colorectal cancer, breast cancer, castrate-resistant prostate cancer, etc. Interestingly, the reduction of H3K36me2 led to increased chemosensitivity of oxaliplatin and 5-fluorouracil in CRC cells [23]. Furthermore, NSD1 depletion has been reported with antitumor efficacy [17] but so far, no reports are available to exploit the NSD1 for improving chemosensitivity and antitumor efficacy.

## Materials and Methods

### Enzymatic Activity Assay

NSD1 enzyme activity was assessed using the NSD1 direct activity chemiluminescent assay kit (BPS Biosciences, USA) is designed to measure NSD1 activity for screening applications. The key to the NSD1 chemiluminescent activity assay kit is a highly specific antibody that recognizes the methylated residue of Histone H3. With this kit, only three simple steps are required for methyltransferase detection. First, S-adenosyl methionine (SAM) is incubated with a sample containing assay buffer and methyltransferase enzyme. Next, a primary antibody is added. Finally, the plate is treated with an HRP-labeled secondary antibody followed by the addition of the HRP substrate to produce chemiluminescence that can then be measured using a chemiluminescence reader. The known methyltransferase inhibitor SAC was identified and profiled for selectivity against the histone methyltransferases NSD1. HTS using the luminescent NSD1 assay has the potential to deliver selective NSD1 inhibitors that may serve as leads in the development of targeted therapies for cancers overexpressing NSD1 [24].

### Cell Culture

DU-145, and HepG2 were obtained from NCCS Pune, India. RWPE1 (normal prostate epithelial cells) was obtained from ATCC (Manassas, VA, USA). Cells were grown under standard conditions of 5% CO_2_ and 37^0^C temperature in a controlled humidified incubator. They were cultured in Dulbecco’s Modified Eagle’s Medium/DMEM (Sigma Aldrich) supplemented with 10% FBS (GIBCO, U.S.A.) and 1% penicillin-streptomycin. RWPE1 was cultured in Keratinocyte Serum Free Media (KSFM) supplemented with 50 mg/L BPE, 5μg/L EGF, and 1% penicillin-streptomycin obtained from ATCC (Manassas, VA, USA). All the cells were used before passage 20. Cells were routinely passage using 0.25% trypsin/0.1% EDTA. Unless otherwise mentioned, all other chemicals were purchased from Sigma (St. Louis, MO, USA).

### Cell viability assay

Cytotoxicity (MTT) assay for 5-O-sulfamoyladenosine (5-SA) was performed in DU145, HePG2, and RWPE1 cells according to the method as described [25]. Cells were harvested, counted, and seeded at a density of 5000 cells per well in a 96-well plate and then incubated for 24 h. Then cells were treated with 5-SA and SAC of various concentrations ranging from 10 µM to 250 µM for 48 h. 2.5 mg of MTT (3- [4, 5-dimethylthiazol-2-yl]-2,5-diphenyltetrazolium bromide) was dissolved in 500 µL of phosphate-buffered saline (PBS) and diluted to 5 mL with serum-free DMEM medium. After 48 h of 5-SA and S-Adenosyl-L-Cysteine treatment (Sigma, St. Louis, MO, USA), 200μl of the MTT solution was added to individual wells. The plate was then wrapped in aluminum foil and incubated at 37 °C for 4 h. The solution in each well, containing media, unbound MTT, and dead cells, was removed by suction. 200μl of DMSO was added to each well. The plate was then kept for shaking and then absorbance was measured at a dual wavelength of 550 nm and 630 nm using a multi-mode automated microplate reader (Flex station III, Molecular Devices, Sunnyvale, CA, USA). The results were expressed as percentage cell viability, assuming the viability of control cells as 100%. Three independent experiments were performed for each study, and all measurements were performed in triplicate.

### Total RNA isolation

Briefly, TRIZOL reagent (Invitrogen, CA, USA) was used for total RNA extraction from cells followed by purification according to the manufacturer’s protocol using RNeasy kit (AuPrep RNeasy Mini Kit; Life Technologies Pvt. Ltd., India). A Nanodrop spectrophotometer (ND-1000) was used to evaluate each sample’s RNA quality and integrity by A260/280 absorbance ratio and agarose gel electrophoresis, respectively [26].

### Reverse transcription and RT-PCR

To evaluate the mRNA (transcript) levels of the NSD1 gene in 5-SA treated samples, a verso cDNA synthesis kit was used for cDNA synthesis from RNA and then quantitative RT-PCR was performed using 2μl of total RNA template (5ng/μl), 2μl of 5X reaction buffer, 1μl of enzyme mix and finally adjusted to 10μl reaction volume using nuclease-free water. The process of Reverse transcription was performed at 42°C for 60 minutes, followed by inactivation at 95°C for 5 minutes. For RT-PCR, 20μl reactions were prepared in the following proportions: 10μl of SYBR Green master mix, 2μl of primer mix, and 8μl of cDNA template (1:80 dilution of the synthesized cDNA in nuclease-free water). RT-PCR was conducted at 95°C for 10 minutes, followed by 40 cycles of 95°C for 10 seconds/60°C for 1 minute using a Light Cycler 2.0 (Roche Diagnostics, USA) [26]. After amplification, melting curve analysis to verify the specificity of the reaction was carried out. Relative changes in NSD1 mRNA expression were assessed using the comparative Ct (ΔCt) method and normalization was done using 18S rRNA as a reference gene. The list of forward and reverse primer sequences of NSD1 and 18S rRNA genes used for mRNA quantification through quantitative RT-PCR is given in Supplementary Table 1.

### Isolation of total proteins and western blotting

Briefly, cells were seeded and allowed to grow till 70-80% confluency in 8 cell culture dishes (2 x 10^6^ cells per plate) in their respective media having 10% FBS and antibiotics (penicillin, 100 IU/ml and streptomycin, 100 μg/ml) under controlled and standard conditions of 5% CO_2_ and 37°C in a well humidified atmosphere of incubator. Afterwards, washing of cells with ice-cold PBS followed by cell lysis in modified low salt buffer (LSB; 10 mM Tris–HCl, pH 7.4, 150 mM EDTA, 1 mM sodium orthovanadate, 10 mM sodium fluoride, 10 mM sodium pyrophosphate, 1 mM phenylmethylsulfonyl fluoride, 10 μg/ml aprotinin, 1 mM sodium butyrate and 0.05% NP-40). It was followed by sonication first and then centrifuged at 12000 g to separate supernatant for the estimation of total cellular proteins and then reduced by using 1X Laemmli’s sample buffer according to a method described by Tikoo et al [26]. Equal samples were run on SDS PAGE and then transferred onto nitrocellulose membrane electrophoretically using semi-dry transfer apparatus (Bio-Rad). Depending on the molecular weight of desired proteins, samples were resolved using sodium dodecyl sulfate-polyacrylamide gels. The membrane-transferred proteins were blocked in 3% BSA for 2 to 3 hours. Further, the membrane was probed with primary and secondary antibodies, developed using HRP chemiluminescent substrate solution, and developed/fixed onto X-ray film. Immunoblotting was performed using an anti-H3K36me2 antibody, anti-NSD1and anti-tubulin antibody (rabbit, 1:1000, Santa Cruz Biotechnology, CA, USA). Incubation of the membrane with horseradish peroxidase (HRP)-coupled secondary antibodies (Santa Cruz Biotechnology, CA, USA) is used to help detect antigen-primary antibody complexes. Densitometry scanning was employed to quantify using NIH Image J software.

### Annexin V/propidium iodide (PI) binding assay

To assess the mode of cell death by 5-SA, 5-FU, or their combination, the Tali® Apoptosis kit

- Annexin V Alexa Fluor® 488 and propidium iodide (Invitrogen) was used according to the manufacturer’s instructions. In short, after the treatment, the cells were collected from 6-well plates using trypsin-EDTA solution, centrifuged at 300 × g for 8 min, resuspended in ABB (Annexin V binding buffer) and incubated with Annexin V Alexa Fluor 488 at room temperature in the dark for 20 min. Following the centrifugation at 300 × g for 5 min, the cells were again resuspended in ABB and incubated with propidium iodide at room temperature in the dark for 5 min. The cells were examined using Tali® Image-Based Cytometer (Invitrogen).

The data were analyzed by FCS Express Research Edition software (version 4.03; De Novo Software, New Jersey, NJ, USA) and expressed as the percentage of cells in each population.

For Flow cytometry-based cell cycle analysis, cells were treated with 5-SA, 5-FU, and 5SAC+5-FU for 48 h. Then, cells were washed with PBS, treated with trypsin-EDTA (Lonza AG, Verviers, Belgium) and fixed with 70% ethanol at 220uC, and conserved at 4uC. For analysis, cells were washed twice with PBS and resuspended in 20 mg/ml RNase (Invitrogen, Cergy-Pontoise, France) and 50 mg/ml propidium iodide (IP) (Invitrogen) to label the DNA. DNA content was measured using PI staining by flow cytometry in BD FACS [27].

### Monolayer wound healing assay

DU145 cells were allowed to grow in 6-well plates supplemented with DMEM media to reach 70–80% confluency as described above. The cells were then incubated in a reduced serum medium for 6 h containing 1% FBS. Cell cultures were scratched with a 200 µL sterile pipette tip and washed with PBS to remove detached cells and debris. Two crosses were scratched in each well, and the scratches were immediately subjected to photography using a Nikon Inverted Microscope at 20 × magnification. Cells were then treated with 5-SA (150μM) and 5-FU (25μM). After 48 h, images of the same areas were acquired by using a Nikon Inverted Microscope at 20 × magnification. The quantitative values of the percentage of cells covered in the scratch after 48 h were determined by using the web-based Wim-Scratch module of Wimasis online software as described [28]. At least three biological replicates per experiment were used and the presented results were representative of triplicate experiments with similar outcomes.

### Cell migration

DU145 cells (5 × 10^4^cells/well) were treated with 5-O-sulfamoyladenosine (150μM) and 5-FU (25μM). After 48 h, cells suspended in 300μl of serum-free DMEM medium were seeded into the upper chamber of each insert (24-well insert; pore size, 8μM; BD Biosciences). Afterward, 500μl of DMEM with 10 % FBS was added to each well of a 24-well plate. The inserts were incubated at 37 °C for 12 h in wells containing DMEM media supplemented with serum. After incubation, migrated cells were washed thoroughly with DPBS, fixed (100 % methanol), followed by staining with crystal violet solution (0.5 % crystal violet in 25 % methanol/DPBS) for 15 min. Cells that didn’t migrate to the lower compartment of inserts were removed with a cotton swab. Photographs of each insert were taken in five random fields (40×). Quantification was expressed in terms of the percentage of area covered with migrated cells using ImageJ software (National Institute of Health, USA).

### Chemosensitivity assay

DU145 cells seeded in 24 well plates (1 x 10^5^ cells per well) were allowed to grow in DMEM medium as described above. Cells were incubated further in serum-free media for 1h and then treated with 5-O-sulfamoyl adenosine (150μM) and 5-FU (25μM). After 48 h, media was removed, PBS washing was carried out, and cell viability was finally assessed.

### In vivo xenograft tumor model

The animal protocol was approved by the Institutional Animal Ethics Committee (IAEC), NIPER (Protocol Approval No. IAEC 20/24). All experiments were performed as per the guidelines of the Committee for Control and Supervision of Experiments on Animals (CPCSEA), India, and NIH guidelines (Guide for the care and use of laboratory animals). Male athymic nude mice (Crl: NU-Foxn1nu) aged 5–6 weeks and weighing 18–22g were obtained from Vivo Bio Tech Ltd, Hyderabad, and transferred to the pathogen-free environment of the National Toxicology Center (NTC) in NIPER. The animals were maintained in sterile and clean cages with HEPA filters under a standard diet with free access to water and controlled conditions of temperature: 20 ± 1 °C, humidity: 50±10%; and 12 h light/dark cycle. All the animals were acclimatized for a period of one week prior to the start of experiments and were properly maintained by taking all necessary precautions and care.

### Injection of DU145 cells into nude mice

Cell derived xenografts (CDX) were developed by subcutaneous injection of viable DU145 (5 × 10^6^) cells dispersed in 1:1 (v/v) of PBS and Matrigel into the flank on the right side of the nude mice [29,30]. After 4 weeks when we observed the animals had attained a tumor volume of 100 mm^3^, we randomly divided them into 5 groups with each group consisting of five animals (n = 5). These 5 groups include cancer control, low dose 5-O-sulfamoyladenosine (LD = 5 mg/kg), high dose 5-O-sulfamoyladenosine (HD = 10 mg/kg), 5-Fluorouracil (5-FU;10 mg/kg) as a positive control group, 5-FU +high dose 5-O-sulfamoyladenosine administered intraperitoneally twice a week for one month before euthanization of mice. Comparable doses of a structurally similar compound (NSC-750854) which are in fact 5-O-sulfamoyladenosine without amino group at position 6 in the adenine group have been already tested in animal xenograft models [22]. This was similar for all groups including the control group. Tumors were measured twice a week with digital calipers, and tumor volume was calculated using the formula: volume = (length × width²)/2, where length represents the longest dimension.

### Histopathological analysis of tumors

A tumor was removed from each animal, sliced, and fixed in 10% v/v formal saline. Thereafter, the tumors were subsequently embedded in paraffin; sections of 5 µm were prepared and mounted on slides previously coated with poly L-lysine. Sections were stained with hematoxylin & eosin to observe the structural changes as described. A cover slip was mounted using DPX and observed at 20×, and 40× magnification using an OLYMPUS BX51 microscope, and the images were captured with an OLYMPUS DP 72 camera attached to the microscope [26].

### Statistical analysis

All the values were expressed as mean ± S.E.M. Statistical comparison between two groups was done using t-test and comparison between more than two different groups was performed using one-way analysis of variance (ANOVA) followed by Tukey’s test. A p-value less than 0.05 was considered as significant.

## Results

### 5-O-sulfamoyladenosine (5-SA) and S-Adenosyl-L-Cysteine (SAC) inhibit NSD1 enzyme activity

In our previous study, 5-O-sulfamoyladenosine (5-SA) was selected after the process of structure-based virtual screening against NSD1, where it appeared as one of the top hit compounds. 5-SA had a high combine score of 2.181 (docking score, XP Gscore, PhaseScreenScore, and MMGBSA delta G Bind) and a leading interaction profile with the active site residues of NSD1better than the endogenous cofactor S-Adenosyl-L-Methionine (SAM), S-Adenosyl-L-Cysteine (SAC) and Sinefungin. SAM is an endogenous cofactor to an active site of NSD1for methyl group transfer. SAC represents a structural or competitive analog of SAM. The chemical structures of SAM, SAC, and 5-SA are shown in Figure 1A. For screening of 5-SA and SAC, NSD1 enzyme activity was assessed using the NSD1 chemiluminescent assay and was analyzed for NSD1 enzyme inhibition potential. Both SAC and 5-SA inhibited NSD1 enzyme activity with their IC_50_ values of 115.003 µM and 53.819 µM respectively (Figure 1B).

**Figure 1.**
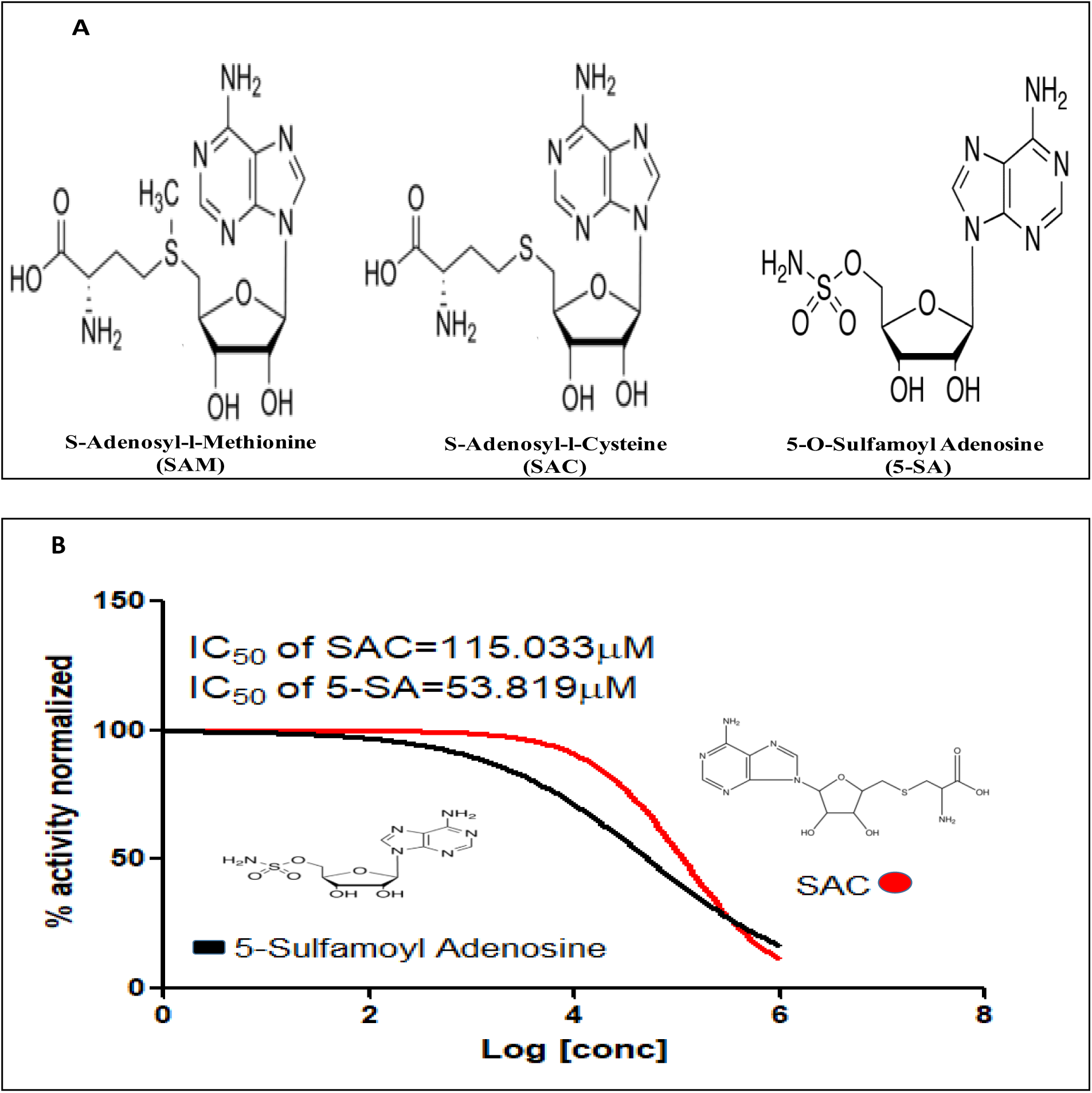
**A)** Chemical structures of relevant compounds in the assay: SAM is an endogenous ligand to the active site of NSD1 for methyl group transfer; S-Adenosyl-L-Cysteine (SAC) as an active set competitive SAM analogue lacking methyl group; 5-SA acting as a competitive inhibitor of SAM as shown in NSD1 chemiluminescent enzyme assay. **B)** S-Adenosyl-L-Cysteine (SAC) and 5-O-sulfamoyladenosine (5-SA) inhibit NSD1 enzyme activity and their respective IC50 values are 115.003 µM and 53.819 µM, respectively.

### 5-SA decreased the viability of DU145 cells, altered NSD1/ H3K36me2 level,and modulated cell sensitivity towards 5-FU

To characterize the effects of 5-SA on cell growth, DU145 cells were treated with 5-SA at different doses in the range of 10 µM to 250 µM for 48 hours. A significant dose-dependent decrease in the growth of cells was observed in comparison to untreated control cells. The IC_50_ of 5-SA in DU145 cells for 48h was calculated as 198 µM (Figure2A). Interestingly, 5-SA and SAC didn’t affect the viability of immortalized human non-tumorigenic prostate epithelial (RWPE1) cells in doses up to 200µM (Supplementary Figure 3A-B). However, 5-SA was relatively more active and inhibited the growth of tumor cells better than the reference compound SAC, which is a SAM analog. The IC_50_ of SAC in DU145 cells for 48h was calculated as 244 µM (Supplementary Figure 3C). The increased expression of NSD1 in prostate cancer (PRAD), liver cancer (LIHC) was evaluated using Cbioportal TCGA datasets (Supplementary Figure 1A, 1B) and strong correlation of increased NSD1 mRNA and its copy number as shown in Supplementary Figure 2A in several cancers, 2B in prostate and liver cancer relevant to our study.

Histone modifications play a crucial role in cell cycle regulation [31]. 5-SA significantly altered the expression of dimethyl levels at H3K36 (H3K36me2) in a dose-dependent manner in DU145 cells (Figure 2B-C). Furthermore, 5-SA significantly altered the expression of NSD1 at protein and mRNA levels in DU145 cells as shown in Figures 2C-F respectively when compared to the untreated control cells. Since epigenetic modulator drugs can modulate the sensitivity of cancer cells to chemotherapeutic agents, we checked whether epigenetic modulator-5-SA treatment sensitizes DU145 cells to 5-fluorouracil. To check the effect of epigenetic modulation by 5-SA on the sensitivity of DU145 (Figure 2G) cells towards 5-Fluorouracil, cells were treated with 5-SA, 5-FU, and the combination of both for 48 h. A significant decrease in cell viability was observed in the combination group of 5-SA and 5-FU due to enhanced cell sensitivity. 5-SA significantly sensitized the tumor cells to 5-FU (around 63% cell death in the combination group vs DU145 control).

**Figure 2.**
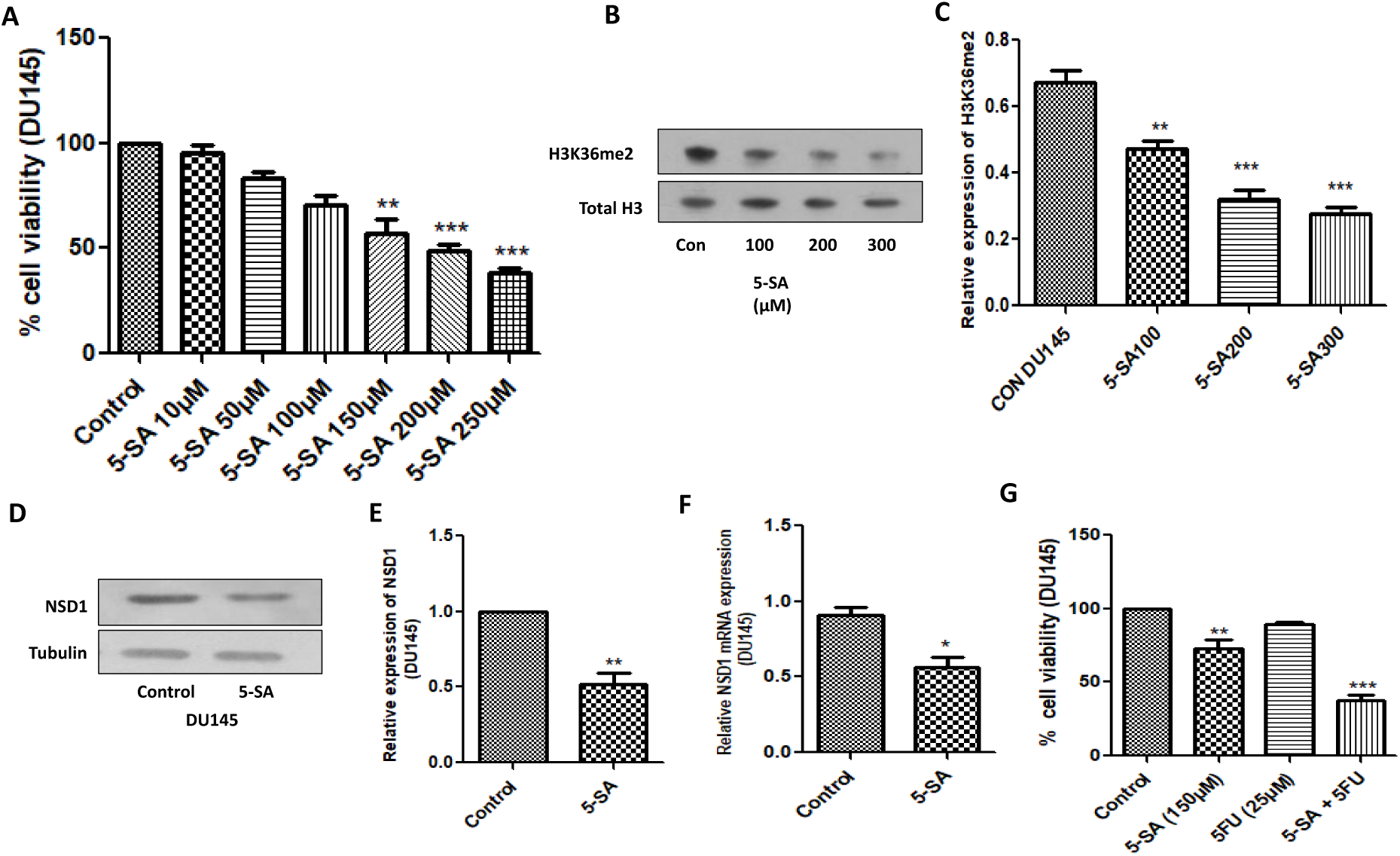
(A)MTT assay andgraphpadtool quantitative analysis of viabilityof cells after 48 h treatment with 5-SA with different concentrations in comparison to control DU145 cells, (B, C, D, E) Western blots and densitometry analysis of H3K36me2 and NSD1 in DU145cells treated with 5-SA. (F) represents the NSD1 mRNA relative changes in 5-SA treated DU145 by RT-PCR and (F) MTT assay of co-treatment with 5-SA and 5-FU for 48 h and graph pad tool quantitative analysis of viability of cells after 48 h treatment with 5-SA (150 µM), 5-FU (25 µM) and the combination in comparison to control DU145cells. Results shown were representativeof threedifferent experiments. All valueswere expressed as mean ± S.E.M. *p < 0.05, **p < 0.01, ***p<0.001 significant vs control group.

### 5-SA and 5-FU impeded the migration and invasion of DU145 cells

To evaluate the impact of 5-SA (150 µM), 5-FU (25 µM), or their combinations on the migration of DU145 cells, a wound healing (scratch) assay was performed. Interestingly, 5-SA significantly inhibited tumor cell migration when compared to the untreated control cells. Wim Scratch tool quantitative analysis revealed that around 81% of the scratch area was covered with cells in the control group, while 60.5%, 68.3%, and 50.1% of the scratch area was covered with cells in the 5-SA, 5-FU, and 5-SA+5-FU treated groups, respectively (Figure 3A). Furthermore, a trans-well migration assay was performed to evaluate the effect of 5-SA (150 µM) and its combination with 5-FU (25 µM) in DU145 cells after 48 h treatment with 5-SA in comparison to DU145 control. 5-SA, 5-FU, and 5-SA+5-FU significantly reduced the invasion of the cells through the trans-well membrane (Figure 3B). However, the impeding effect of the combination was significantly more than the other two individual treatments.

**Figure 3.**
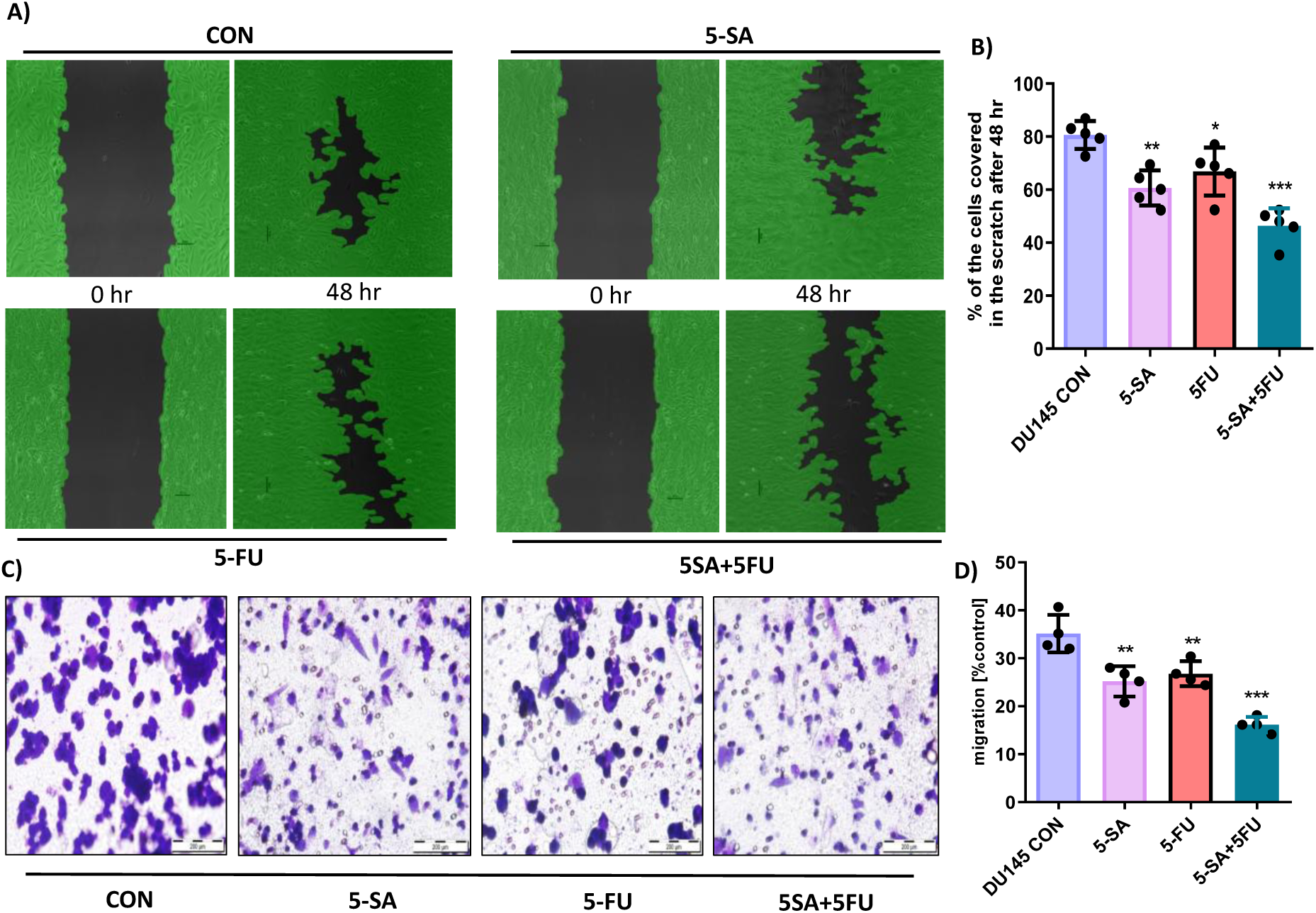
5-SA and 5-FU impeded themigrationand invasionof DU145 cells. (A) Scratch wound images and Wim Scratch tool quantitative analysis of the percentage of the cells covered in the scratch after 48 h treatment with 5-SA (150 µM) and 5-FU (25 µM) in comparison to DU145 control. (B) Transwell migration assay with 5-SA and 5-FU in DU145 cells after 48 h treatment with 5-SA (150 µM) and 5-FU (25 µM) in comparison to DU145 control. The results shown were representative of three different experiments. All values were expressed as mean ± S.E.M. **p<0.01 significant vs control group ***p<0.001 significant vs control group.

### 5-SA induced apoptosis and modulated the cell cycle phase of DU145-treated cells in the presence of 5-FU

Treatment of DU145 cells with 5-SA (150 µM), 5-FU (25 µM), or their combination induced apoptosis. The Annexin V/propidium iodide (PI) binding assay results showed that exposure of DU145 cells to 5-SA and 5-FU and their combination led to a significant increase of apoptosis in all three groups relative to the untreated control. Quantitatively, the % apoptosis was 37%, 28% and 57% in the 5-SA group, 5-FU group and 5-SA+5-FU combination group respectively (Figure 4A). In addition to this, the % dead cells were 2%, 24% and 9% in the respective groups. Similarly, the percentage of live cells was 60%, 49% and 34% respectively as shown in Supplementary Figure 4A.

**Figure 4.**
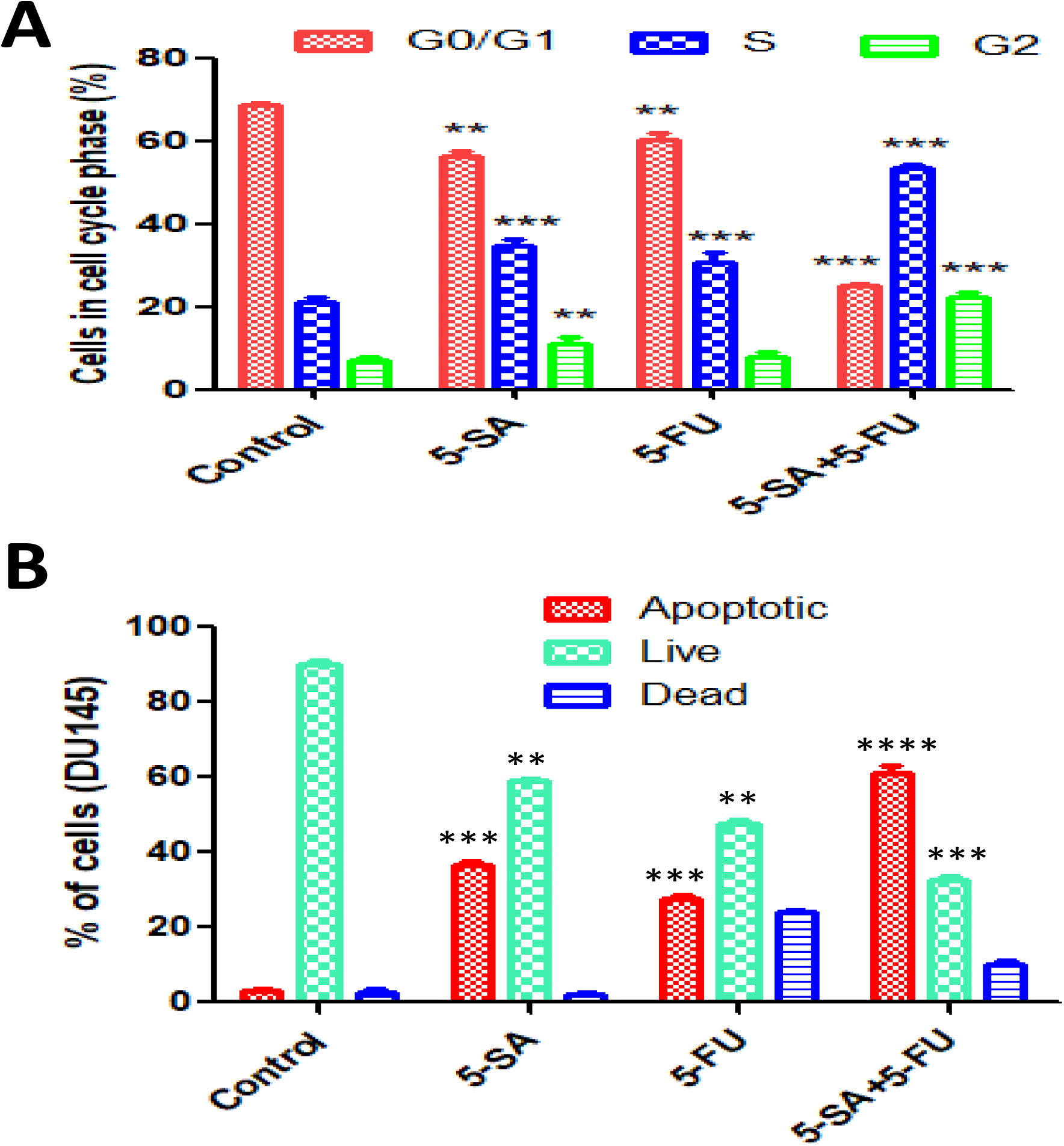
Effect of 5-SA and 5-FU on cell cycle profile and apoptosis of DU145 cells. (A) Apoptosis assay using Tali image-based cytometer quantified in DU145 cells, control, and others treated with 5-SA (150 µM) and 5-FU (25 µM) for 48 h in comparison to DU145 control. (B) Cell cycle analysis was performed using PI staining by flow cytometry in BD FACS Verse flow cytometer. Representative cell cycle histogram of the DU145 cells, control, and others treated with 5-SA (150 µM), 5-FU (25 µM), and 5-SA+5-FU for 48 h. The bar diagram compares variations in cell distribution percentagein eachphase of the cell cycle of DU145 cells after treatments as compared to control. 5-SA and 5-FU and their combination induced apoptosis in DU145 cells. Data represented as mean ± SEM of independent experiments conducted in triplicates. ANOVA followed by Turkey’s multiple comparison tests was performed to test the significance of the data. **p < 0.01, ***p < 0.001 vs. control.

The effects of 5-SA, 5-FU, or their combination on the cell cycle profile of DU145 prostate cancer cells were assessed. The cell cycle distribution analysis showed that the exposure of DU145 cells to both 5-SA and 5-FU in combination led to a significant increase of cells at the S phase (P < 0.05, P < 0.01), while there was a significant decrease at the G0/G1 phase (P < 0.01) (Figure 4B). The percentage of DU145 cells at the S phase increased from 20.3 ± 1.099% (control) to 32.9 ± 1.64%, 28.2 ± 2.5% and 54.4 ± 1.05%, after treatment with 150μM 5-SA, 25μM 5-FU and a combination of both for 48 h respectively as shown in Supplementary Figure 4B. The percentage of DU145 cells at the G0/G1 phase decreased from 71.0 ± 0.349% (control) to 57.5 ± 1.049%, 61.8 ± 1.54%, 24.5 ± 0.4% in 5-SA, 5-FU and 5SA+5-FU treated groups respectively. However, cells at the G2/M phase were not significantly changed after the respective treatment (P < 0.05).

### 5-SA decreased the viability of HepG2 cells, altered NSD1 expression and improved sensitivity towards 5-FU

To re-evaluate the effect of 5-SA on the cell viability and NSD1 expression level, we chose to treat another cell line HepG2 due to the significant expression of NSD1 [17]. In line with the above results, we find that the effects of 5-SA on the growth of HepG2 cells at different doses in the range of 10 µM to 250 µM for 48 hours were significant. A dose-dependent decrease in the viability of HepG2 cells was observed compared to the positive control (untreated cells). The IC_50_value of 5-SA in HepG2 cells for 48 h was calculated as 168.3 µM (Figure 5A). To check the effect of epigenetic modulation by 5-SA on the sensitivity of HepG2 cells towards 5-Fluorouracil, cells were treated with 5-SA, 5-FU, and the combination of both for 48 h (Figure 5B). A significant decrease in cell viability was observed in the combination group of 5-SA and 5-FU due to enhanced cell sensitivity. 5-SA significantly sensitized the tumor cells to 5-FU (around 72% cell death in the combination group vs HepG2 control. Furthermore, 5-SA significantly altered the expression of NSD1 at mRNA and protein levels in HepG2 cells as shown in Figures 5C and 5D, respectively when compared to the untreated control cells. However, 5-SA was relatively more active and inhibited the growth of tumor cells better than the SAC, which is a SAM analog. The IC_50_ of SAC in HepG2 cells for 48h was calculated as 281.9 µM (Supplementary Figure 3D).

**Figure 5.**
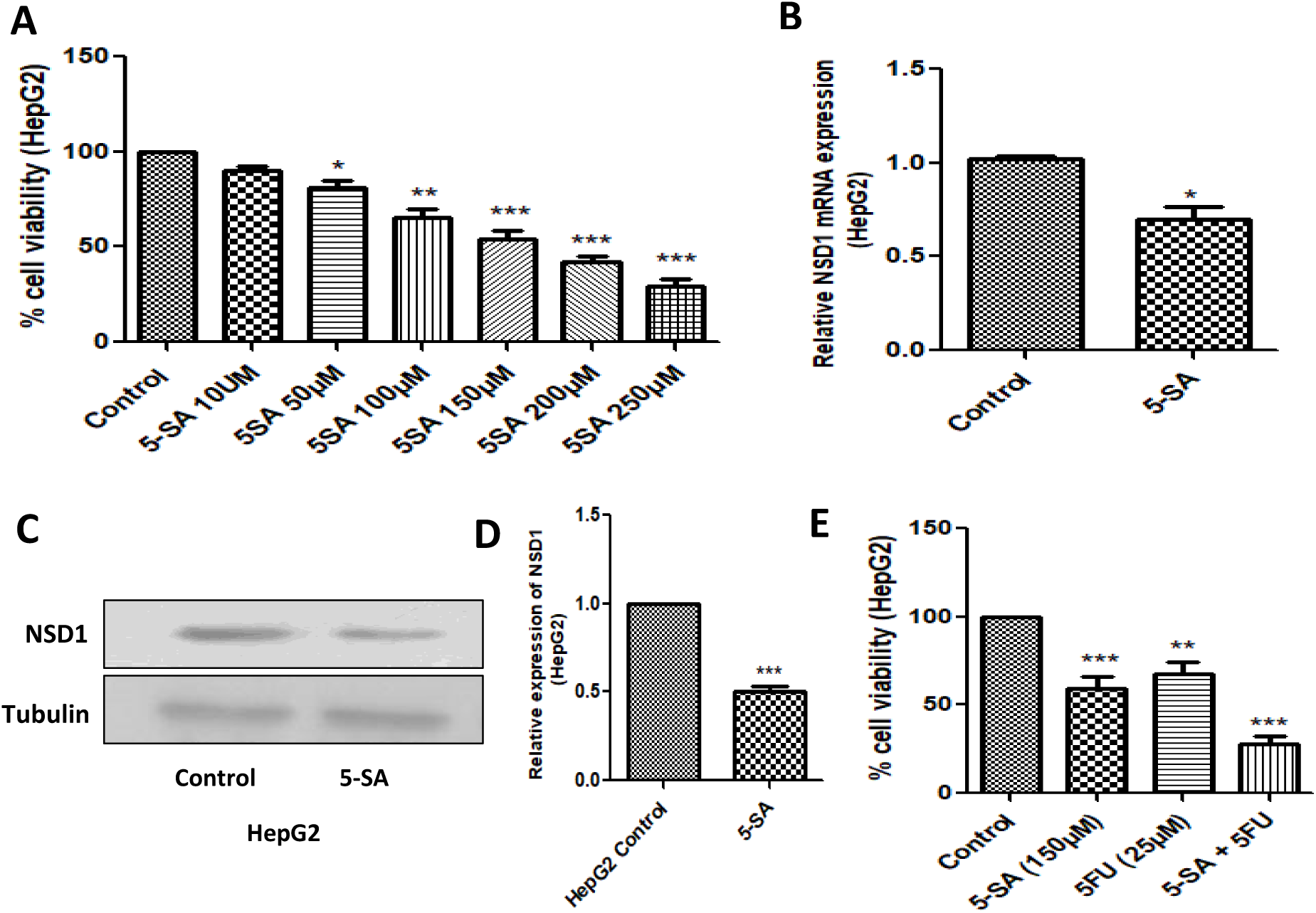
(A)MTT assay andgraphpadtool quantitative analysis of viabilityof cells after 48 h treatment with 5-SA with different concentrations in comparison to control in HepG2 cells. (B) Represents the NSD1 mRNA relative changes in 5-SA treated DU145 by RT-PCR in comparison to control HepG2 cells. (C) Western blots and densitometry analysis of NSD1 in DU145 treated cells with 5-SA. (D) MTT assay of co-treatment with 5-SA and 5-FU for 48 h and graph pad tool quantitative analysis of viability of cells after 48 h treatment with 5-SA (150 µM), 5-FU (25 µM) and combination in comparison to control HepG2cells The results shown were representative of three different experiments. All values were expressed as mean ± S.E.M. *p < 0.05, **p < 0.01, ***p<0.001 significant vs control group.

### 5-SA, 5-FU, and their combination showed anti-cancer effects in the DU145-induced prostate cancer xenograft model

We observed significant anticancer activity of 5SA, 5-FU, and combination in DU145 cells (in-vitro). Further, the evaluation of the antitumor efficacy of 5-SA *in vivo* requires the development of a xenograft animal model for prostate cancer using DU145 cells. To ensure whether 5-SA, 5-FU or their combination also shows good anti-cancer effects *in vivo*, we developed a xenograft model of prostate cancer, by subcutaneous injection of DU145 cells into the right flank of nude mice and evaluated its efficacy (Supplementary Figure 5). Tumors were allowed to grow to around 100 mm^3^ tumor volume and thereafter, the animals were divided into five groups (n = 5). Control mice received 0.9% saline solution. The tumor volume was checked twice weekly for up to four weeks. 5-fluorouracil (5-FU) was used as positive control in our study. 5-FU alone caused suppression of tumor volume to some extent. High dose 5-SA significantly decreased the size of the tumors when compared to the control group. 5-SA + 5-FU combination exerted the maximum effect on the tumor regression as evident from Figures 6 and 7. Schematic representation showing xenograft tumor growth, drug treatment, and surgical resection of tumors (Figure 6A). Representative macroscopic images of sacrificed mice and surgically removed tumors are shown in Figure 7A. The respective treatment groups showed cytotoxic effects on the tumor as evidenced by the H&E staining (Figure 7B). The combination group of 5-SA and 5-FU showed marked cytotoxic damage. Moreover, Figure 6 indicates that the percent tumor growth inhibition was highest, and average tumor weights were lower in the 5-SA +5-FU group than in the other respective groups. Percentage change in body weight was assessed in all animals of each group to determine the toxicological effects of each treatment. However, the % change in body weight was significant in 5-SA -HD, 5-FU, and the combination treatment group (5-SA +5-FU). In addition, the Kaplan Meier survival analysis indicated that the 5-SA, 5-FU, and their combination improved the survival of the mice (Figure 6C). However, survival was much improved in the combination group of 5-SA +5-FU than in the individual treatment groups possibly because of improved chemosensitivity.

**Figure 6.**
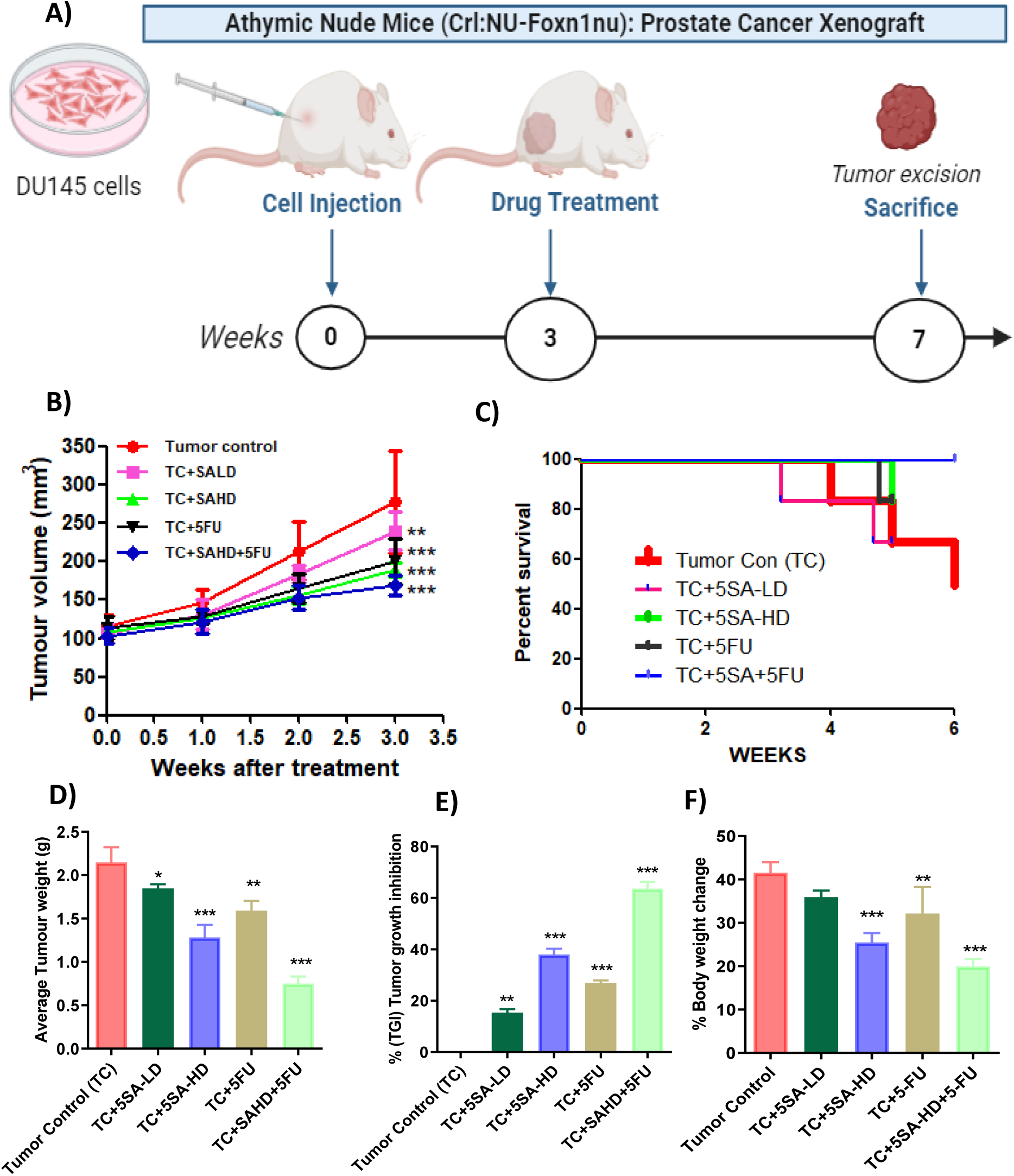
Effect of 5-SA and 5-FU treatment on the A) Schematic representation of tumor implantation and drug treatment, B) tumor volume-time curve, C) Kaplan-Meier survival curve D) average tumor weight), % tumor growth inhibition (TGI), and F) % body weight change. All values were expressed as mean ± S.E.M. *p<0.05, **p<0.01, ***p<0.001 significant vs tumor control (TC) group.

**Figure 7.**
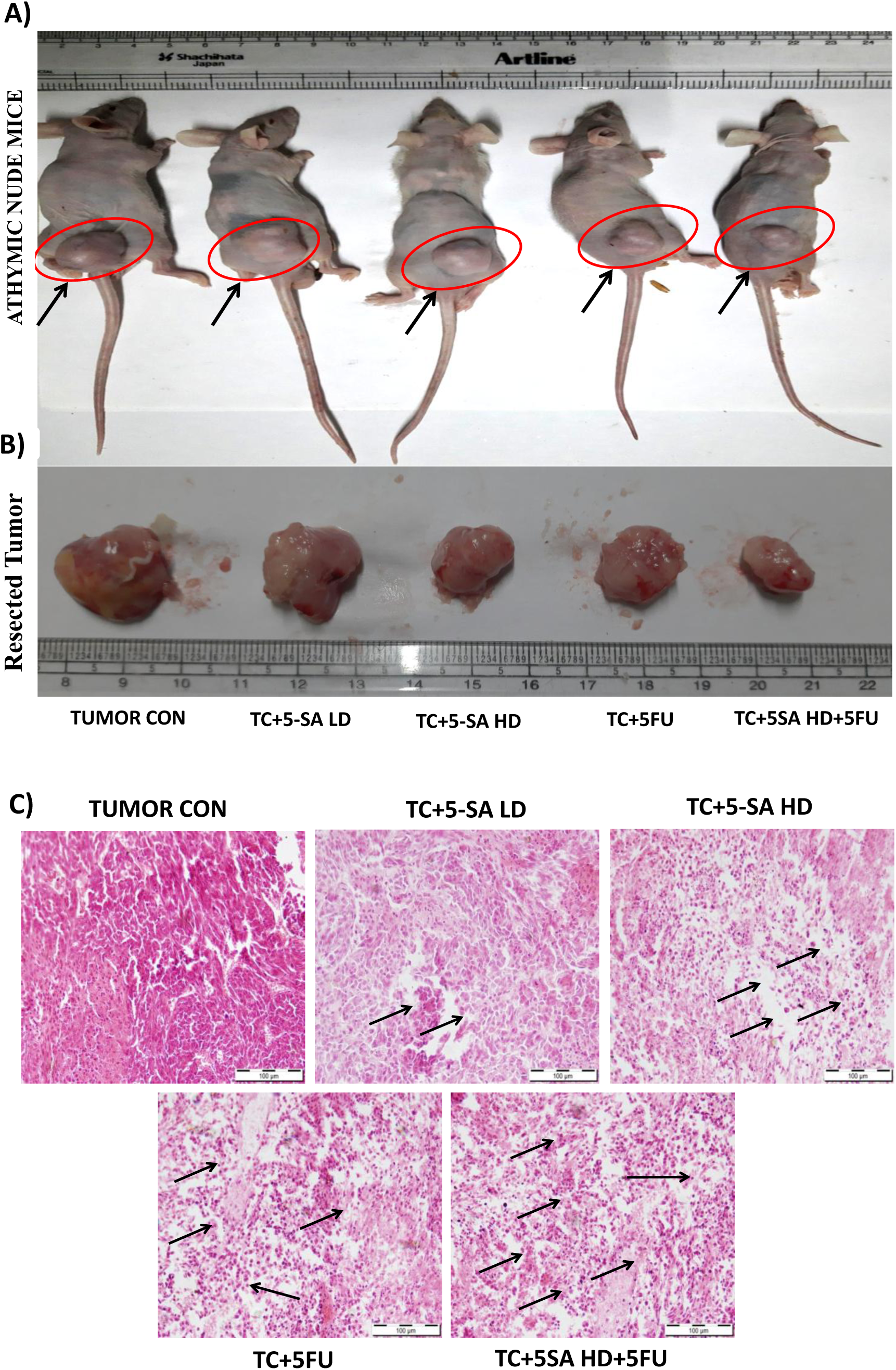
Effect of 5-SA and 5-FU treatment on the tumor size in DU145-induced prostate cancer xenograft. (A) Representative macroscopic images depicting tumor growth of xenograft nude mice from different groups. Images of sacrificed animals before surgical tumor removal of mice in each group (n = 5), and images of surgically removed tumors of mice in each group. B) Histopathological analysis of tumors excised from tumor control (TC), 5-SA (LD = 5mg/kg, 5-SA (HD = 10mg/kg), 10 mg/kg 5-fluorouracil (5-FU) and 5-fluorouracil + 5-SA (HD) treated animals. The arrows depict the interstitial spaces in tumor samples. No such significant spaces were found in tumors of untreated animals.

## Discussion

Emerging studies suggest that nuclear receptor-binding SET domain 1 (NSD1) is elevated in multiple cancers such as acute myeloid leukemia (AML) [8,13], multiple myeloma (MM) [9], lung cancer [14], metastatic prostate cancer [15], metastatic pancreatic duct adenocarcinoma (PDA) [16] and hepatocellular carcinoma (HCC) [17]. In addition, TCGA clinical data analysis using Cbioportal also revealed the overexpression of NSD1 in prostate and hepatocellular cancer (Supplementary Figures 1 and 2). We have recently reported that structure-based virtual screening of large chemical libraries (ZINC and ChemDiv) against NSD1 identified 5-SA as one of the potent inhibitors for NSD1 [12]. In this study, we first evaluated enzyme inhibition involving SAM (endogenous ligand for NSD1) as a natural substrate for methyl transfer while SAC and 5-SA as competitive analogs lacking methyl group, inhibited NSD1 enzyme activity with IC_50_ values of 115.003 µM and 53.819 µM, respectively. Sinefungin, a known NSD inhibitor, inhibits NSD1 with IC_50_ value of 46±1 µM [32] and suppressed invasion of hepatoma cell line C1-30 at 400 µM [33]. Treatment of Sinefungin for 48 hr. decreased the viability of Hela, K462, and MCF7 cells with IC_50_ value of around 100 µM [34]. However, Sinefungin is shown to be nephrotoxic [35]. Drug toxicity and ADME properties are essential parameters for the safety profile and pharmacokinetics of drug molecules. Further, various properties of 5-SA, SAC, and Sinefungin predicted and compared using the Lazar toxicity predictor [36] and Swiss ADME predictor [37]. All the compounds have favorable toxicity/ safety profiles and other properties (water-solubility, lipophilicity, and pharmacokinetics). However, the non-carcinogenic value of 5-SA (0.319) was relatively on the better side than that of Sinefungin (0.281) and SAC (0.301) as indicated in Supplementary Table 2.

5-SA significantly decreased the viability of tumorigenic DU145 and HepG2 cells in a dose-dependent manner with IC_50_ values of 198 µM and 168 µM respectively, while no such effect was observed in non-tumor prostate epithelial RWPE1 cells. The study highlights that DZNep as an epigenetic modulator of enzyme EZH2 inhibit tumor cells while having a slight effect on non-tumor cells [38]. NSD1 is primarily known to regulate gene expression through methylation of lysine 36 on histone H3 (H3K36) [4,5]. Treatment of 5-SA significantly reduced the expression of H3K36me2 in a dose-dependent manner in DU145 cells compared to the control. 5-SA significantly decreased the protein and mRNA expression of NSD1 in DU145 and HepG2 cells. Mechanistically, NSD1 knockout in mouse embryonic stem cells (mESCs) led to the decrease of H3K36me2 at active enhancers [39].

5-SA impeded the migration and invasion of DU145 cells. Interestingly, 5-SA also inhibited tumor cell proliferation and migration as compared to control. Wim-Scratch tool quantitative analysis revealed that the maximum area of the scratch was covered with cells in the untreated control group and the minimum scratch area was covered in treated groups, particularly in the combination group (5-SA +5-FU). Additionally, the transwell migration assay showed that 5-SA and its combination with 5-FU significantly reduced the migration of cells through the transwell membrane compared to untreated DU145 cells. However, the impeding effect of the combination was significantly greater than individual treatments. NSD1 knockout suppresses hepatocellular carcinoma cell proliferation and migration [17]. Several reports suggest that epigenetic modulator drugs can modulate the sensitivity of cancer cells to chemotherapeutic agents [40–42]. Moreover, Suramin with multiple mechanisms of action including epigenetic modulation of NSD enzymes has also shown chemosensitivity potential with 5-FU and other agents in several studies and clinical trials [43,44]. In line with this observation, we found that 5-SA also increased the sensitivity of DU145 and HepG2 cells to the cytotoxic effects of 5-FU in a Chemosensitivity assay.

Histone modifications play a crucial role in cell cycle regulation [45]. The combination of 5-SA +5-FU significantly altered the cell cycle phase with respect to the control cells due to the increased accumulation of cells in the S phase and the corresponding decrease in the G0/G1 phase. Studies reveal that 5-FU treatment of oral cancer cells increased in S-phase cells [46]. Epigenetic modulators may alter the histone modifications and influence the cell cycle process and apoptosis. It has also been reported that silencing of NSD1 promotes cell apoptosis in association with inflammation [47].

In support of *in vitro* studies, the antitumor efficacy of 5-SA *in vivo* using a xenograft model of prostate cancer induced by DU145 cells revealed that 5-SA alone significantly suppressed tumor volume. Due to chemo-sensitization and enhanced cytotoxic damage to the tumor microenvironment as evident from H&E staining, 5-SA -HD+5-FU markedly reduced tumor weight, volume, and % body weight change compared to control and other individual treatment groups. Previous studies have also shown that increased interstitial space amplified penetration of paclitaxel/ doxorubicin and subsequently induced more apoptosis in cancers [48,49]. In our study, the enhanced tumor reduction in the 5-SA + 5-FU group might be due to enhanced penetration due to increased interstitial spaces. Similarly, the % tumor growth inhibition (% TGI) was significantly improved in all treatment groups compared to the control group and was highest in the combination group. Anticancer agents and combination therapies that target tumor microenvironment prolong survival and initiate marked inhibition of tumor growth [50–52]. Interestingly, the Kaplan Meier survival analysis revealed that the combination of 5-SA with 5-FU markedly improved the survival of the mice followed by the survival status with 5-SA and 5-FU respectively. Thus, epigenetic modulation using 5-SA to alter NSD1 activity can be of profound clinical significance, as it provides a novel therapeutic approach to sensitize tumors towards chemotherapeutic agent 5-FU. However, further studies are required to extrapolate the role and efficacy of 5-SA with other chemotherapeutic agents and in other types of cancers. Additionally, our study provides a pharmacologically distinct platform to exploit novel opportunities in designing more potent NSD1 inhibitors.

## Author Contributions

Zahid Rafiq designed and performed *in vitro* and *in vivo* experimentation and wrote the manuscript. Kulbhushan Tikoo designed the overall concept of the study, supervised the experiments, and approved the final version of the manuscript.

## Acknowledgement

This work was supported by grants from the National Institute of Pharmaceutical Education and Research, S.A.S. Nagar, India, and the Indian Council of Medical Education and Research (ICMR) in New Delhi, India.

## Data availability

All data included in the text, figures, and supporting information are provided.

## Declarations

### Ethics approval and consent to participate

All animal use procedures were conducted in strict accordance with the guidelines of the Committee for Control and Supervision of Experiments on Animals (CPCSEA), India, and NIH guidelines (Guide for the care and use of laboratory animals). The study was approved by the Institutional Animal Ethics Committee (IAEC), NIPER (Protocol Approval No. IAEC 20/24).

