## Supplementary Information for "Inhibition of NSD1 by 5-O-Sulfamoyl Adenosine improved 5-FU sensitivity by suppressing cancer cell proliferation and xenograft tumor growth"

[1] Laboratory of Epigenetics and Diseases, Department of Pharmacology and Toxicology, National Institute of Pharmaceutical Education and Research (NIPER) S.A.S. Nagar, India

[*]Corresponding author: Prof. Kulbhushan Tikoo, Department of Pharmacology and Toxicology, National Institute of Pharmaceutical Education and Research (NIPER) S.A.S. Nagar, India,

*This work was supported by grants from the National Institute of Pharmaceutical Education and Research, S.A.S. Nagar, India*

***Supplementary Figure 1.*** *NSD1 mRNA expression based on HiSeq_RNASeqV2 TCGA utilizing Cbioportal, (A) LIHC, and (B) PRAD datasets showing the gene overexpression in terms of amplification gain and diploid functions.*

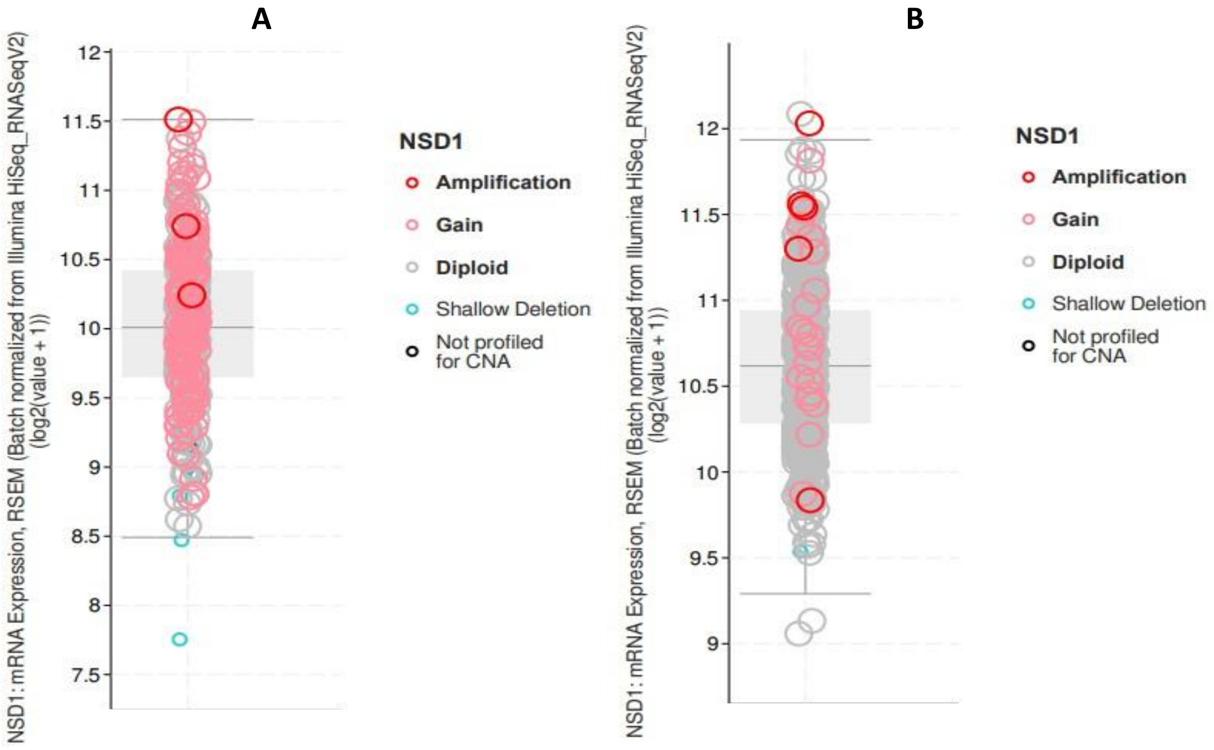


**LIHC-**
**Liver Hepatocellular Carcinoma**
**(TCGA, Pan Cancer Atlas)**

**PRAD-**
**Prostate Adenocarcinoma**
**(TCGA, Pan Cancer Atlas)**

***Supplementary Figure 2.*** *The expression of NSD1 in 38 cancer cell lines was obtained from the cancer cell line encyclopedia showing the strong correlation of increased NSD1 mRNA and its copy number, A) in several cancers, B) in prostate and liver cancer as indicated by the arrows in blue relevant to our study.*

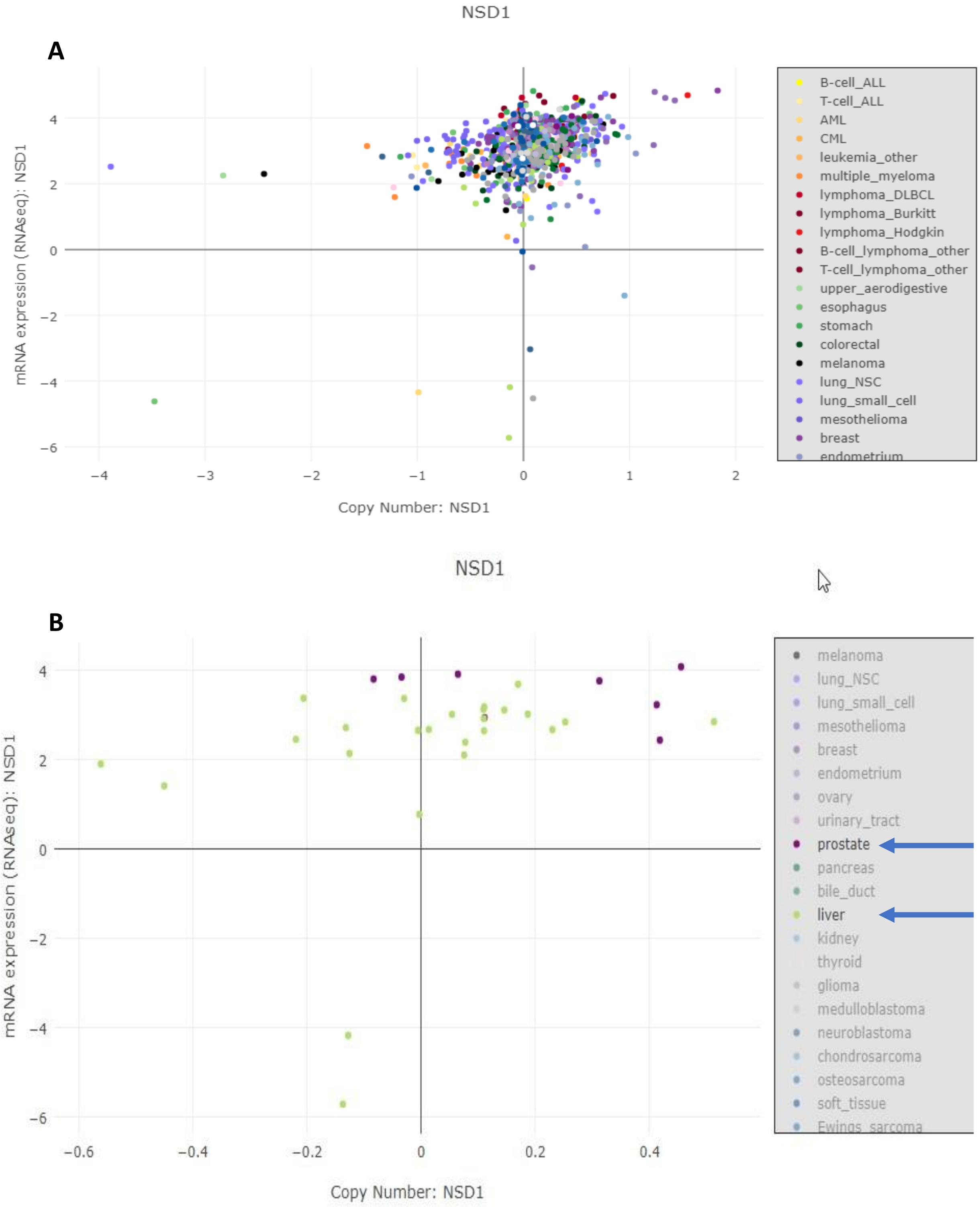

***Supplementary Figure 3.*** *MTT assay and graph pad tool quantitative analysis of the viability of benign RWPE1 prostate epithelial cells and tumorigenic DU145 and HepG2 cells after 48 h treatment with 5-SA and* SAC *with different concentrations in comparison to (A, B) control RWPE1, (C) control DU145 cell, and (D) control HepG2 cells.*

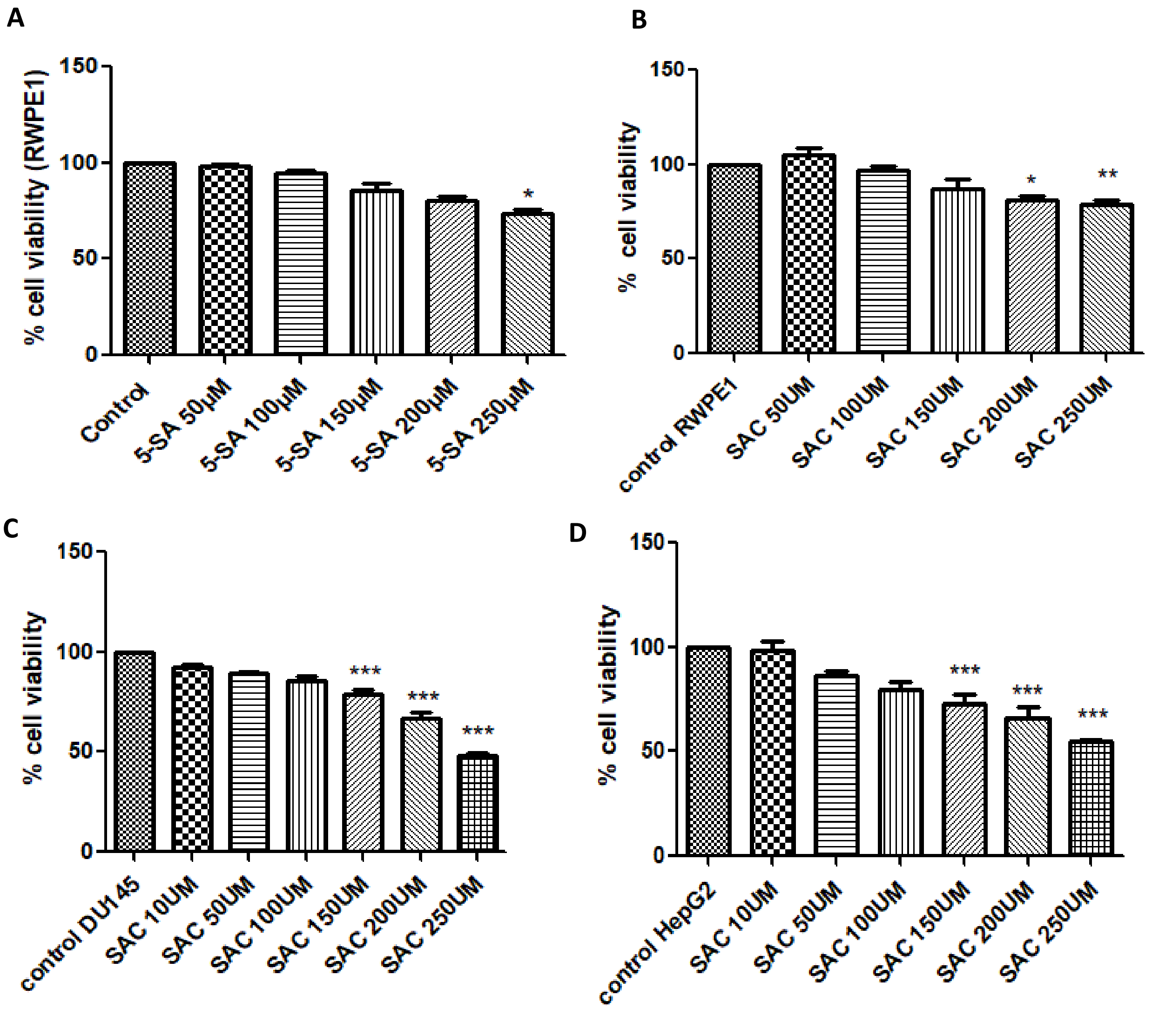


To characterize the effects of 5-SA and SAC, which is a competitive SAM analog on cell growth, non-tumorigenic benign RWPE1 and tumorigenic DU145 and HEPG2 cells were treated with 5-SA and SAC at different doses ranging from 10 µM or 50 µM to 250 µM for a period of 48 hours. A significant dose-dependent decrease in the viability of cells was observed in comparison to untreated cell control. The $IC_{50}$ values in DU145 and HepG2 cell lines for 48 hr. were calculated as 244 µM (Figure 2A) and 281.9 µM (Figure 3C-D) respectively.

***Supplementary Table 1.*** *Sequence of primers used for mRNA quantitation by RT-PCR.*

| S.No | GENE NAME | PRIMER SEQUENCE |
|---|---|---|
| 1. | NSD1 | Fwd: CAAGGAAGCGAAAACGACAGAGG<br><br>Rev: CCGTCCTGTGAGGCATTAGTTC |
| 2. | 18S rRNA | Fwd: GCAATTATTCCCCATGAACG<br><br>Rev: AGGGCCTCACTAAACCATCC |

***Supplementary Table 2.*** *Profiling of compound 5-SA and SAC against NSD2 and NSD3 isoforms*

| Isoforms | Inhibitory activity ($IC_{50}$ ± SD) in µM. | |
|---|---|---|
| | **5-SA** | **SAC** |
| NSD1 | 53.819± 0.32 | 115.013± 0.32 |
| NSD2 | 150.94 ± 0.38 | 103.20 ± 0.56 |
| NSD3 | 1.39 ± 0.56 | 160.43 ± 0.53 |

*Supplementary Figure 4. 5-SA induced apoptosis and modulated the cell cycle phase of DU145-treated cells in the presence of 5-FU.*

### I) Effect of 5-SA and 5-FU on the apoptosis of DU145 cells

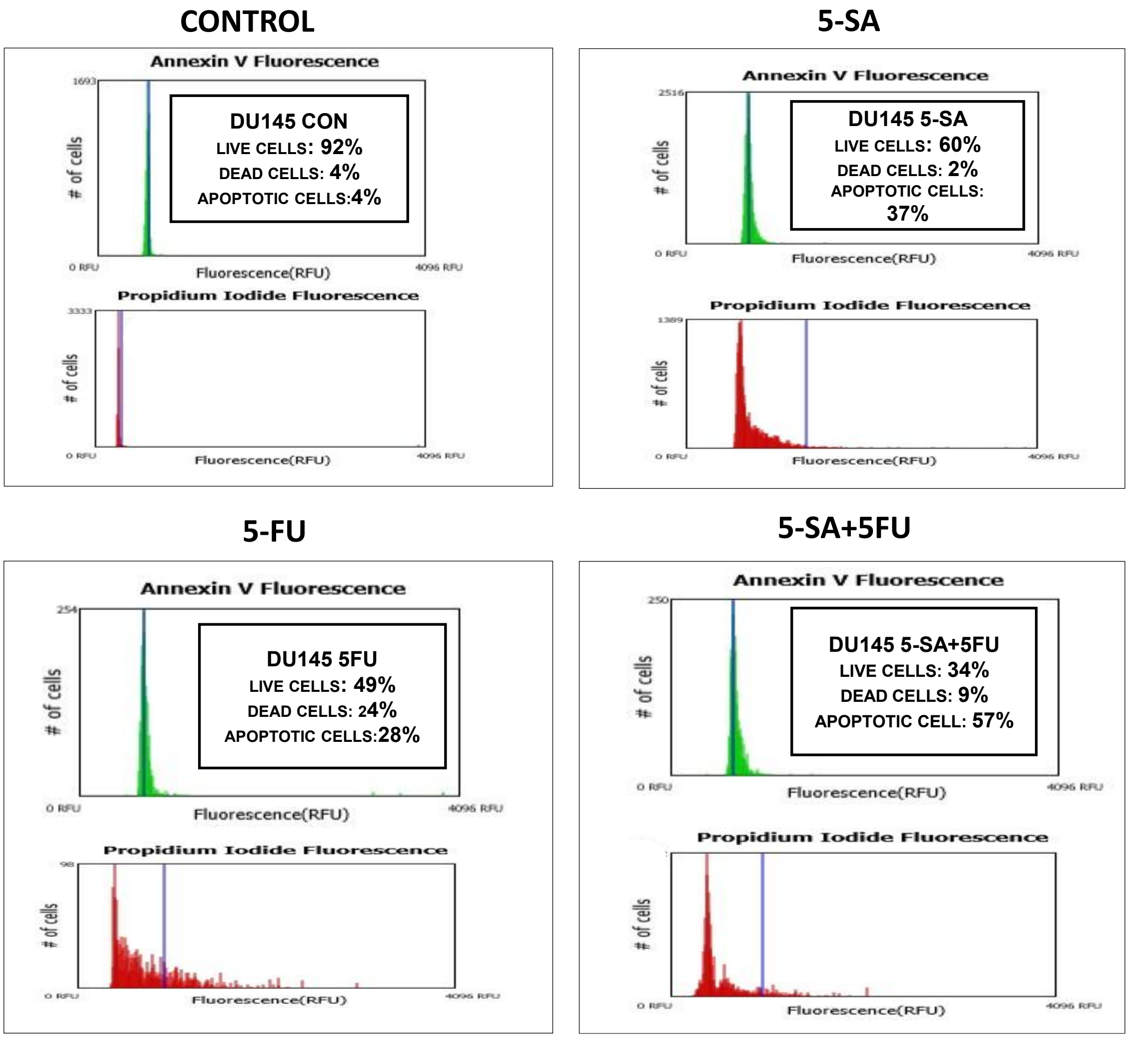

### II) Effect of 5-SA and 5-FU on Cell Cycle analysis of DU145 cells

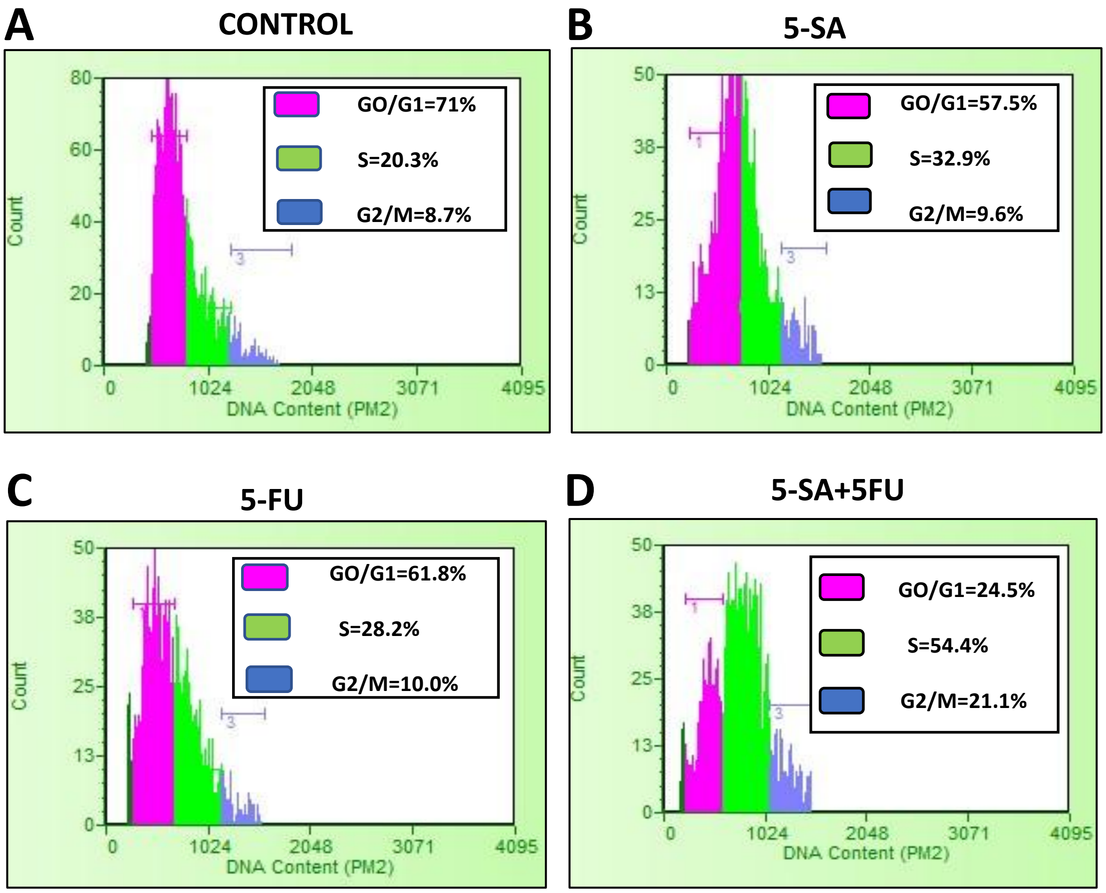

***Supplementary Figure 5.*** *Diagrammatic representation of tumor model development in nude mice DU145 induced xenograft.*

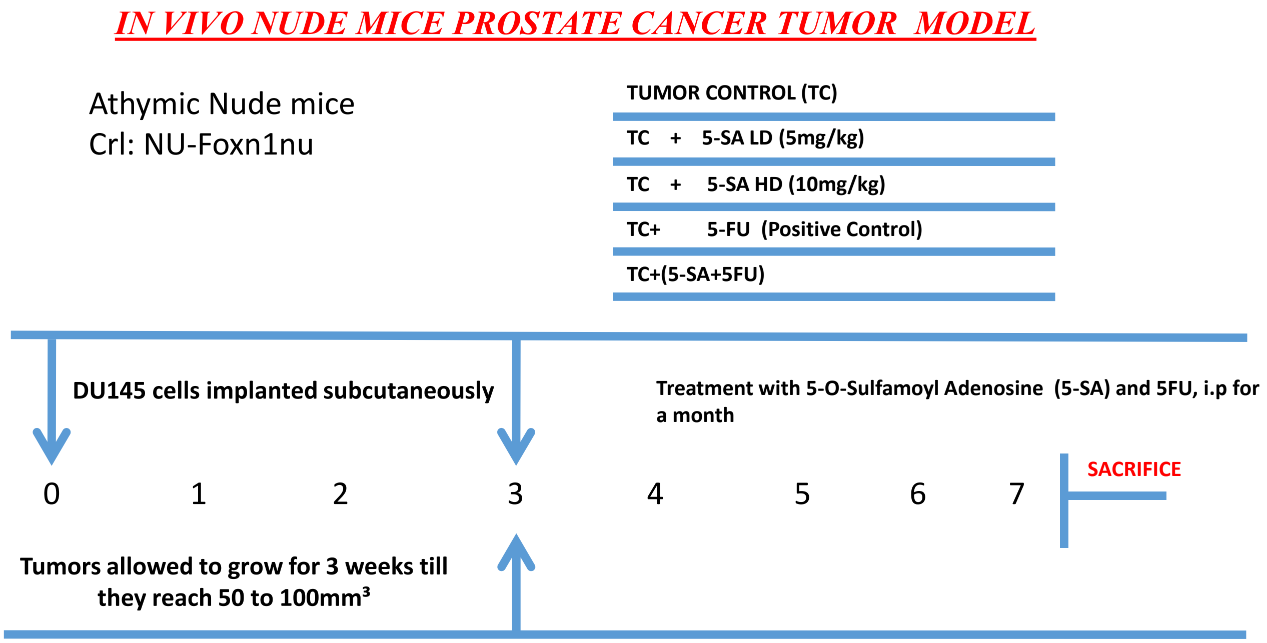

***Supplementary Table 2.*** *Prediction of in-silico toxicity by Lazar toxicity, and ADME properties of 5-SA, SAC, and Sinefungin (SIN).*

| | Toxicity prediction | | In silico ADME | | |
|---|---|---|---|---|---|
| **S.NO** | **Carcinogenicity (RODENTS)** | **Mutagenicity (Salmonella typhimurium)** | **Physicochemical properties** | **Water solubility** | **Pharmacokinetics** |
| **5SA** | **Prediction : non-carcinogenic**<br>**Probability :**<br>**non-carcinogenic : 0.319** | **Prediction: non-mutagenic** | **Formula: C10H14N6O6S**<br>**Molecular weight: 346.32g/mol**<br>**Molar refractivity: 74.27**<br>**TPSA: 197.08Å²** | **Log S (SILICOS-IT): 0.56**<br>**Solubility: 1.27e+03 mg/ml; 3.67e+00 mol/l**<br>**Class: soluble** | **BBB permeant NO**<br>**P-gp substrate NO**<br>**CYP1A2 inhibitor NO**<br>**CYP2C19 inhibitor NO**<br>**CYP2C9 inhibitor NO**<br>**CYP2D6 inhibitor NO**<br>**CYP3A4 inhibitor NO** |
| **SAC** | **Prediction: non-carcinogenic**<br>**Probability:**<br>**noncarcinogenic: 0.301** | **Prediction: non-mutagenic** | **Formula: C13H18N6O5S**<br>**Molecular weight: 370.38 g/mol**<br>**Molar refractivity: 88.00**<br>**TPSA: 207.93 Å²** | **Log S (SILICOS-IT): 0.31**<br>**Solubility: 7.56e+02 mg/ml; 2.04e+00 mol/l**<br>**Class: soluble** | **BBB permeant NO**<br>**P-gp substrate NO**<br>**CYP1A2 inhibitor NO**<br>**CYP2C19 inhibitor NO**<br>**CYP2C9 inhibitor NO**<br>**CYP2D6 inhibitor NO**<br>**CYP3A4 inhibitor NO** |
| **SIN** | **Prediction: non-carcinogenic**<br>**Probability:**<br>**Noncarcinogenic: 0.281** | **Prediction: non-mutagenic** | **Formula: C15H23N7O5**<br>**Molecular weight: 381.39 g/mol**<br>**Molar refractivity: 92.73**<br>**TPSA: 208.65 Å²** | **Log S (SILICOS-IT): 0.34**<br>**Solubility: 8.25e+02 mg/ml; 2.16e+00 mol/l**<br>**Class: soluble** | **BBB permeant NO**<br>**P-gp substrate NO**<br>**CYP1A2 inhibitor NO**<br>**CYP2C19 inhibitor NO**<br>**CYP2C9 inhibitor NO**<br>**CYP2D6 inhibitor NO**<br>**CYP3A4 inhibitor NO** |

Uncropped Images

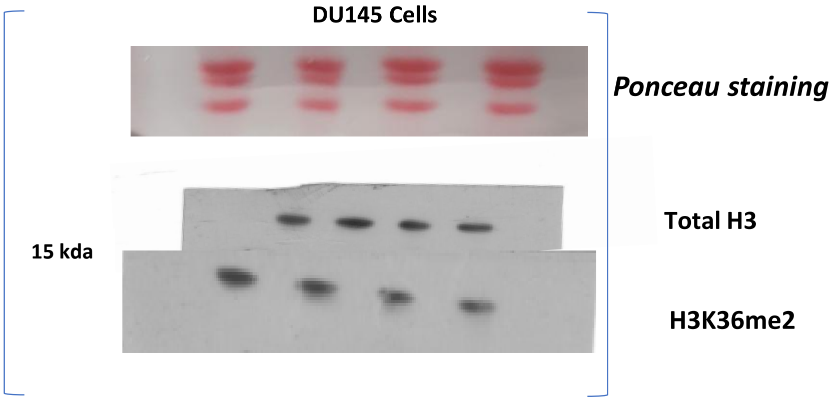


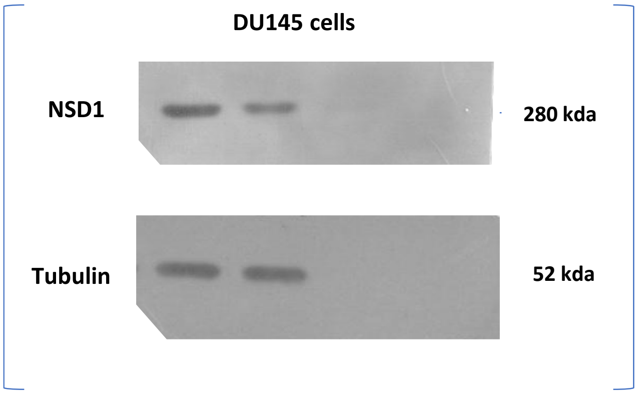


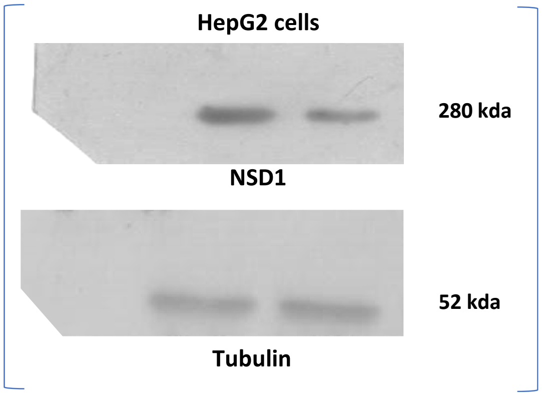
